# Post-mitotic centriole disengagement and maturation leads to centrosome amplification in polyploid trophoblast giant cells

**DOI:** 10.1101/2022.05.25.493455

**Authors:** Garrison Buss, Miranda B. Stratton, Ljiljana Milenkovic, Tim Stearns

**Affiliations:** Department of Molecular and Cellular Physiology, Stanford University School of Medicine, Stanford, CA 94305, USA; Department of Biology, Stanford University, Stanford, CA 94305, USA; Department of Genetics, Stanford University School of Medicine, Stanford, CA 94305, USA

**Author notes:** Authors contributed equally to this work (Garrison Buss, Miranda B. Stratton).

## Abstract

DNA replication is normally coupled with centriole duplication in the cell cycle. Trophoblast giant cells (TGCs) of the placenta undergo endocycles resulting in polyploidy but their centriole state is not known. We used a cell culture model for TGC differentiation to examine centriole and centrosome number and properties. Prior to differentiation, trophoblast stem cells (TSCs) have either two centrioles before duplication, or four centrioles after. We find that average nuclear area increases approximately 8-fold over differentiation, but most TGCs do not have more than four centrioles. However, these centrioles become disengaged, acquire centrosome proteins, and can nucleate microtubules. In addition, some TGCs undergo further duplication and disengagement of centrioles, resulting in substantially higher numbers. Live imaging revealed that disengagement and separation are centriole autonomous and can occur asynchronously. Centriole amplification, when present, occurs by the standard mechanism of one centriole generating one procentriole. PLK4 inihibition blocks centriole formation in differentiating TGCs but does not affect endocycle progression. In summary, centrioles in TGC endocycles undergo disengagement and conversion to centrosomes. This increases centrosome number, but to a limited extent compared with DNA reduplication.

## Introduction

Centrosomes are the major microtubule organizing center (MTOC) in mammalian cells during interphase and assist in spindle formation during mitosis. They consist of two main elements: 1) a pair of centrioles, defined as microtubule-based cylindrical structures that are ∼400 nm in length with nine-fold symmetry, and 2) associated pericentriolar material (PCM) that imparts the centrioles with MTOC activity (Nigg & Holland, 2018; Nigg & Stearns, 2011). Centrosomes are unique in that, along with chromosomal DNA, they are the only structures known to be duplicated exactly once per cell division cycle in animal cells. In mitotic cycles, newly-formed centrioles are engaged to their parental centriole and are embedded in the PCM of the parental centriole. Upon passage through mitosis, the engagement link is broken, and new centrioles undergo the centriole-to-centrosome conversion and acquire their own PCM (W. J. Wang et al., 2011). Disengaged centrioles usually remain close to each other by the action of cohesion fibers, until they separate at the initiation of mitosis (Nigg & Stearns, 2011; G. Wang et al., 2014). The duplication cycle of centrosomes ensures that there are two centrioles in a G1 cell, which duplicate in S-phase and are then segregated as pairs on the mitotic spindle (Tsou & Stearns, 2006; G. Wang et al., 2014). In the context of cells that adhere to this canonical form of centriole duplication and segregation, aberrant centriole number is detrimental (Godinho & Pellman, 2014).

Centrioles are required to form a primary cilium for critical cell signaling pathways. One rationale for the tight control of centriole number is to ensure that only a single cilium can form in most cells (Nigg & Stearns, 2011). Centriole loss both prevents formation of the primary cilium and results in mitotic defects leading to p53-dependent cell cycle arrest (Lambrus et al., 2015). Conversely, having too many centrioles results in the formation of multiple cilia, compromising their function in signal transduction, as well as formation of multipolar spindles, interfering with chromosome segregation (Godinho & Pellman, 2014; Mahjoub & Stearns, 2012). The presence of extra centrioles is a hallmark of many cancers, can itself promote tumorigenesis and invasion, and is strongly prognostic of poor patient outcomes (Basto et al., 2008; Denu et al., 2016).

Although the centriole duplication and DNA replication cycles are coupled in most animal cells, there are examples of differentiated cells that alter their DNA replication, centriole duplication, or cell division cycles to specifically amplify DNA or centrioles. For example, multiciliated cells (MCCs) found in the airway epithelium, brain ependyma, and oviduct, have hundreds of centrioles and associated motile cilia that are used to generate directional fluid flow (Klos Dehring et al., 2013; Spassky & Meunier, 2017; Vladar & Stearns, 2007). MCCs engage a specific transcriptional program during differentiation that results in massive centriole amplification without concomitant DNA replication (Kyrousi et al., 2015; Vanderlaan et al., 1983; Vladar et al., 2018).

There are also many examples in which amplification of DNA content has been observed, largely in the context of cells undergoing endocycles (Edgar et al., 2014; Macauley et al., 1998; Schoenfelder et al., 2014; Ullah et al., 2008). Endocycles encompass a range of cell cycle behaviors. Remarkably, little is known about the coordination of replication of centrioles and DNA in such endocycles. We seek here to determine whether endocycling cells specifically amplify DNA and not centrioles, separating the two cycles as in MCCs, or whether both are coordinately amplified during endocycles.

Mammalian trophoblast giant cells (TGCs) are an important polyploid cell type that establishes the maternal-fetal interface for nutrient, oxygen, and waste exchange (Silva and Serakides, 2016). Both the mother and fetus contribute to the formation, and in turn, the function of the developing placenta (Cross, 2005; Silva & Serakides, 2016; Simmons & Cross, 2005). TGCs invade and remodel the maternal decidua, which is critical for embryonic implantation and placentation (Maltepe & Fisher, 2015). Defects in trophoblast invasion can lead to pregnancy-related disease states; for example, preeclampsia is characterized by a loss or reduction of trophoblastic invasiveness, while gestational trophoblastic disease is characterized by increased invasiveness (Maltepe & Fisher, 2015; Silva & Serakides, 2016). TGCs have been studied as models of polyploidy with respect to genome amplification (Edgar et al., 2014; Ullah et al., 2008). There are several types of TGCs, based on their derivation and/or location in the placenta. These represent different paths towards polyploidy, including endocycling, cell fusion and cell division failure. (Klisch et al., 2017; Sakaue-Sawano et al., 2013; Simmons et al., 2007; Simmons & Cross, 2005; Zybina & Zybina, 2005; Zybina & Zybina, 1996).

To address questions about the coordination of DNA and centriole duplication in endocycles, we focus here on murine mononuclear TGCs, which are derived by differentiation of murine trophoblast stem cells (TSCs). Mononuclear TSCs exit the canonical cell cycle and enter an endocycle in which DNA replication continues up to 64N without an intervening cell division (Ullah et al., 2008). We show that centrioles in endocycling TGCs undergo centriole disengagement and conversion to centrosomes without passage through mitosis. This leads to an increase in centrosome number but only to a limited extent compared with DNA amplification. Thus, TGCs, like multiciliated epithelial cells, are able to uncouple the centriole and DNA cycles as part of a differentiation program.

## Results

### Polyploidization of murine TGCs in the developing placenta and in vitro

To investigate centriole and centrosome number in murine TGCs, we sought to establish conditions for *in vitro* differentiation that would represent those *in vivo*. Figure 1A shows a cartoon representation of the placenta, with the zone containing TGCs outlined in red. Figure 1B shows a section of the mouse conceptus at day 9.5 (e9.5), with the corresponding region containing TGCs, evident by their larger nuclear area. Cells in this zone were imaged in subsequent experiments as comparison to *in vitro*-produced TGCs (see below). We exploited an existing cell culture model to differentiate TGCs from derived trophoblast stem cells (TSCs) *in vitro* (Hannibal & Baker, 2016). Wild-type mouse TSCs isolated from blastocyst embryos were stimulated to differentiate into TGCs by shifting growth medium, as depicted in Figure 1C. Briefly, TSCs were grown in the presence of growth factors activin A, fibroblast growth factor-4 (FGF-4), and heparin. At the beginning of differentiation (t=0d), the growth factors were removed and the culture medium was supplemented with retinoic acid for up to 10 days to encourage adoption of the TGC fate (Simmons et al., 2007). Differentiation was accompanied both by an increase in cell size (Figure 1C) and by induction of the canonical TGC marker placental lactogen-I alpha, *Prl3d1* (Hemberger et al., 2004; Rai & Cross, 2015; Simmons et al., 2007; Simmons & Cross, 2005) (Figure 1D).

**Figure 1:**
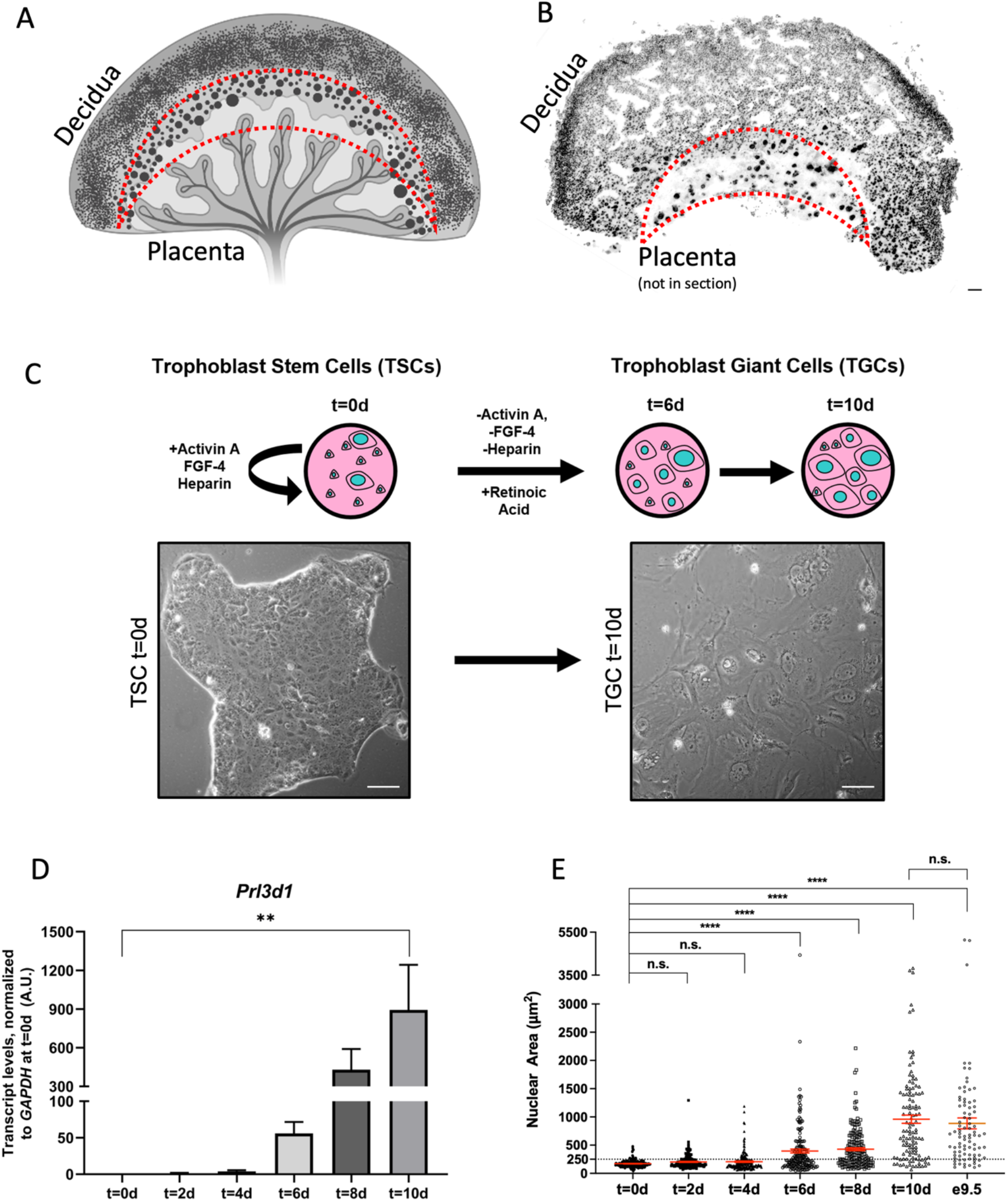
Trophoblast Stem Cells can be propagated and differentiated into TGCs *in vitro:* A. Diagram of the developing mouse placenta (light grey and vasculature) and decidua (dark gray), with the layer of TGCs outlined in red. Graphic was created with Biorender.com. B. Widefield image of a tissue section of conceptus at e9.5 stained with DAPI; layer of TGCs, characterized by large nuclei, is outlined in red. Scale bar = 100 µm C. Schematic of trophoblast giant cell (TGC) differentiation time course for detailed protocol, see Methods. TSCs were differentiated into TGCs for up to 10 days, with a time point collected every two days. The t=0d time point was collected at the beginning of differentiation, thus cells at t=0d are also considered to be TSCs. Note that TGC differentiation is an asynchronous process, and there may be TGCs with large nuclei present at the beginning of differentiation, as indicated in the schematic. Phase images of TSCs (left) and TGCs (right) show clear morphological differences of the two cultures. Scale bar: 50 µm. D. To validate TGC differentiation over the time course, expression of TGC-specific lactogen *Prl3d1* was evaluated by quantitative real time-PCR. Data were collected in four independent experiments. *Gapdh* probe was used to normalize samples, expression is relative to time t=0d **p-value ≤0.01 E. Quantification of nuclear area throughout TGC differentiation shows a gradual increase in size of nuclei. Graph shows average nuclear area (mean plus the standard error of the mean) measured during *in vitro* differentiation time course. Data were collected in three independent experiments. ****p-value ≤0.0001, n.s., not significant.

To assess differentiation at the single-cell level, we chose to use nuclear size as a proxy for TGC differentiation as this correlates with increase in TGC ploidy and DNA content (Morimoto et al., 2021; Roukos et al., 2015; Supplementary Figure 1). Nuclear area enlargement has been previously shown to correlate with the increase in TGC marker gene expression, with most non-expressing cells falling in the range 118-249 µm^2^. (Carney, et al., 1993). Thus, we defined TGCs as mononuclear cells with a nuclear area of ≥ 250 µm^2^ (Figure 1E). At t=0d, 90% of cells had nuclei smaller than this cutoff. At t=10d, 83% of cells had nuclei larger than this cutoff, with a mean of 960 +/- 72 µm^2^. In e9.5 placenta sections, 84% of cells in the TGC zone had nuclei larger than this cutoff, with a mean of 886 +/- 96 µm^2^, suggesting that the TGC in vitro differentiation conditions adequately allow for TGC polyploidization.

These results also show that some TSCs spontaneously differentiate into TGCs in vitro, and that not all cells become TGCs even when stimulated by the described treatments, consistent with previous findings (Yan et al., 2001). This results in a heterogeneous population of trophoblast cells at any given time point, with the fraction of cells adopting the TGC fate increasing over time (Figure 1D,E). We therefore consider a nuclear area of ≥ 250 µm2 and differentiation time t=6d to be the onset of TGC fate for most of the population.

### Centriole and centrosome number increase during TGC endocycles

Chromosomal DNA in mononuclear TGCs is amplified exponentially via serial S-phases without an intervening mitosis (Sakaue-Sawano et al., 2013; Zybina & Zybina, 1996). Thus, a simple hypothesis would be that centriole number would also increase exponentially, in coordination with DNA replication. We first investigated centriole number in TGCs in the natural context of the developing placenta of a mouse strain expressing eGFP-centrin2 to visualize centrioles and Arl13B-mCherry to mark primary cilia in embryonic cells (Bangs et al., 2015). Imaging sections of the TGC-containing zone of the e9.5 placenta showed both cells with normal-sized nuclei and cells with large nuclei and positive Cytokeratin 7 staining, indicative of being TGCs (Maldonado-Estrada et al., 2004). Examples of two fields of cells are shown in Figure 2A. The non-TGCs in these sections usually had two eGFP-centrin2 foci. This is as expected for canonical centriole duplication, where the centrin foci represent either two centrioles pre-duplication, or four centrioles in two engaged pairs post-duplication. TGCs usually had a higher number of eGFP-centrin2 foci, although the large cell size precluded a definitive determination of centriole number per cell (Figure 2A). In no case were primary cilia observed, consistent with previous work (Bangs et al., 2015).

**Figure 2:**
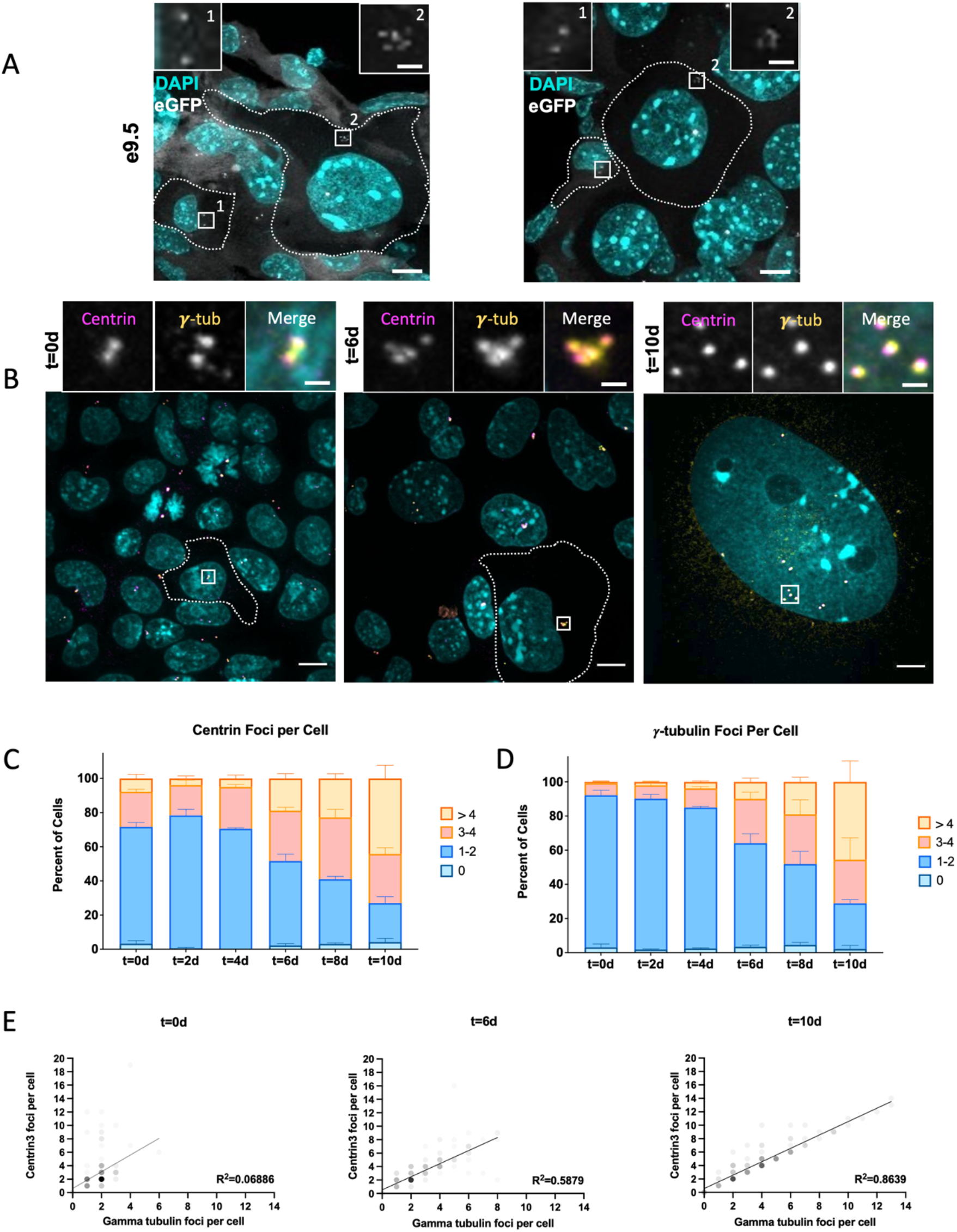
Increase in centriole number and centrosome number during TGC differentiation A. Confocal microscopy images of tissue sections of a mouse conceptus (e9.5) transgenic for *eGFP-centrin2/Arl13b-mCherry*. Sections correspond to the TGC layer described in 1A. DAPI (cyan) was used to visualize nuclei, and the native eGFP-centrin2 fluorescence to visualize centrioles (white). TGCs were identified by nuclear size relative to adjoining cells. Scale bars: overview, 10 µm; inset 2µm. B. Microscopic images of differentiating TGCs at t=0d, t=6d, and t=10d. Cells were fixed and labeled with antibodies to mark the centrosome (***γ***-tubulin, yellow) and centrioles (centrin, magenta). DAPI (cyan) was used to visualize nuclei. Centrioles from cells indicated with dashed lines are shown at higher magnification in insets. The t=10d images are from a large TGC whose boundary is beyond shown field of view. Scale bars: overview, 10 µm; inset 2µm. C. Quantification of centriole number throughout TGC differentiation as measured by centrin immunofluorescence, shows an increase in centriole number as cells differentiate. The percent of cells with the indicated centriole numbers was calculated in three independent experiments. For each experiment, a minimum of 60 cells per condition were counted; bars represent the mean percent of cells. Error bars represent the standard error of the mean (SEM). D. Quantification of centrosome number throughout TGC differentiation time course, as marked by ***γ***-tubulin immunofluorescence. The percent of cells with the indicated centriole numbers was calculated in three independent experiments. For each experiment, a minimum of 60 cells per condition were counted; bars represent the mean percent of cells. Error bars represent the standard error of the mean (SEM). E. Correlation of centriole and centrosome number per cell as identified by centrin and ***γ***-tubulin immunofluorescence, respectively, at the beginning of differentiation (t=0d) middle of differentiation (t=6d) and end of differentiation (t=10d) from the data in C and D, shading opacity of each point is 10%.

We next used the *in vitro* cell culture system to assess centriole and centrosome number during TGC differentiation (Figures 2B-E; Figures S2A-C). Cells were imaged by immunofluorescence staining for centrin and ***γ***-tubulin, a component of the PCM. As expected, all ***γ***-tubulin foci had at least one associated centrin focus. We counted any such ***γ***-tubulin foci as a centrosome and all individually-resolvable centrin foci as centrioles (Figure 2B,C). At t=0d, the majority of TSCs in the population had two centrosomes, each with a single centriole (Figures 2B-D; Figure S2C), typical of diploid G1 cells.

Over the course of 10 days of TGC differentiation, there was an increase in centriole and centrosome number (Figures 2B-D, Figures S2A,B). The fraction of cells with only 2 centrioles decreased, coincident with the increase in cells with 4 centrioles and 4+ centrioles (Figure 2B). Centrosome number increased in a similar manner over the time course, with the percentage of cells having two centrosomes decreasing to become a minority within the population (Figure 2D). Interestingly, over this time course, many TGCs also increased their centriole and centrosome numbers beyond what would be expected for a cycling cell. At t=10d, 40% of TGCs had a higher-than-normal number of centrioles (>4 centrin foci per cell) and 65% had a higher-than-normal number of centrosomes (>2 ***γ***-tubulin foci per cell) (Figure 2B-D).

Although the number of centrioles rose over time in differentiating TGCs, the increase was modest and not a function of 2^n^ (1-2 extra centrioles per cell). Even if the analysis is restricted to cells with very large, single nuclei (≥1000 µm^2^), which presumably had undergone one or more endocycles (DNA ≥8n), the mean centriole number in these cells was 5.2, lower than the expected value of 8. Remarkably, the number of centrosomes, as determined by ***γ***-tubulin foci, was the same in that population, with 5.2 centrosomes/cell.

We observed that the number of centrioles was equal to the number of centrosomes in t=10d TGCs, suggesting that centrioles were not engaged and that they had acquired the PCM component ***γ***-tubulin, despite not having passed through mitosis. This interpretation was consistent with images of t=10d TGCs, showing that each focus of centrin was associated with a focus of ***γ***-tubulin (Figure 2B). These single centrioles in t=10d TGCs were often dispersed, compared to centrioles in TSCs which were usually closely associated, as expected for centrioles linked by cohesion fibers.

Given the marked similarities between the distributions of centrin and ***γ***-tubulin during TGC differentiation, we quantified the centriole/centrosome ratio per cell over time. The number of cells with amplified centrosomes and exactly a 1:1 ratio of centrin to ***γ***-tubulin increased over time to reach ∼40% of all cells in the population by t=10d (Figure S2C). Even when the ratio of centriole to centrosome was not exactly 1:1, there was a clear trend for TGCs to approach that ratio throughout differentiation (Figure 2E). By t=10d, the relationship between centrin and ***γ***-tubulin per cell increased to become strongly correlated, with R^2^ = 0.8639 (Figure 2E). These results show that centrioles in differentiating TGCs typically go through at least one duplication event before becoming disengaged in the absence of mitosis, that these centrioles undergo a conversion from centrioles to centrosomes during the endocycle, and that some TGCs gain supernumerary centrioles and centrosomes during differentiation.

### Centrioles disengage in differentiating post-mitotic TGCs

To determine how new centrioles are formed in differentiating TGCs and how they might become single-centriole centrosomes, we observed centriole dynamics in living cells (Figure 3). For this purpose, we derived TSCs from blastocysts using the eGFP-centrin2; Arl13b-mCherry mouse strain, as previously described (Bangs et al., 2015; Kidder, 2014; Tanaka, 2006). These TSCs were plated on gridded 35 mm imaging dishes and were induced to differentiate as in Figure 1C. Beginning on day four of differentiation, centrioles were imaged by spinning-disk confocal microscopy for at least 48 h. Fields of mononucleate cells that were chosen were at the edge of colonies, such that cells had space to expand or divide over the course of imaging. As expected for the asynchronous differentiation process, these fields contained a mix of TSCs, early TGCs and late TGCs. With a few exceptions for cases of centriole amplification, we focused our analysis on mononuclear cells that had two centrin foci (i.e. either two centrioles or two pairs of centrioles) with the rationale that it would be easiest to observe changes to the normal pattern of duplication in such cells.

**Figure 3:**
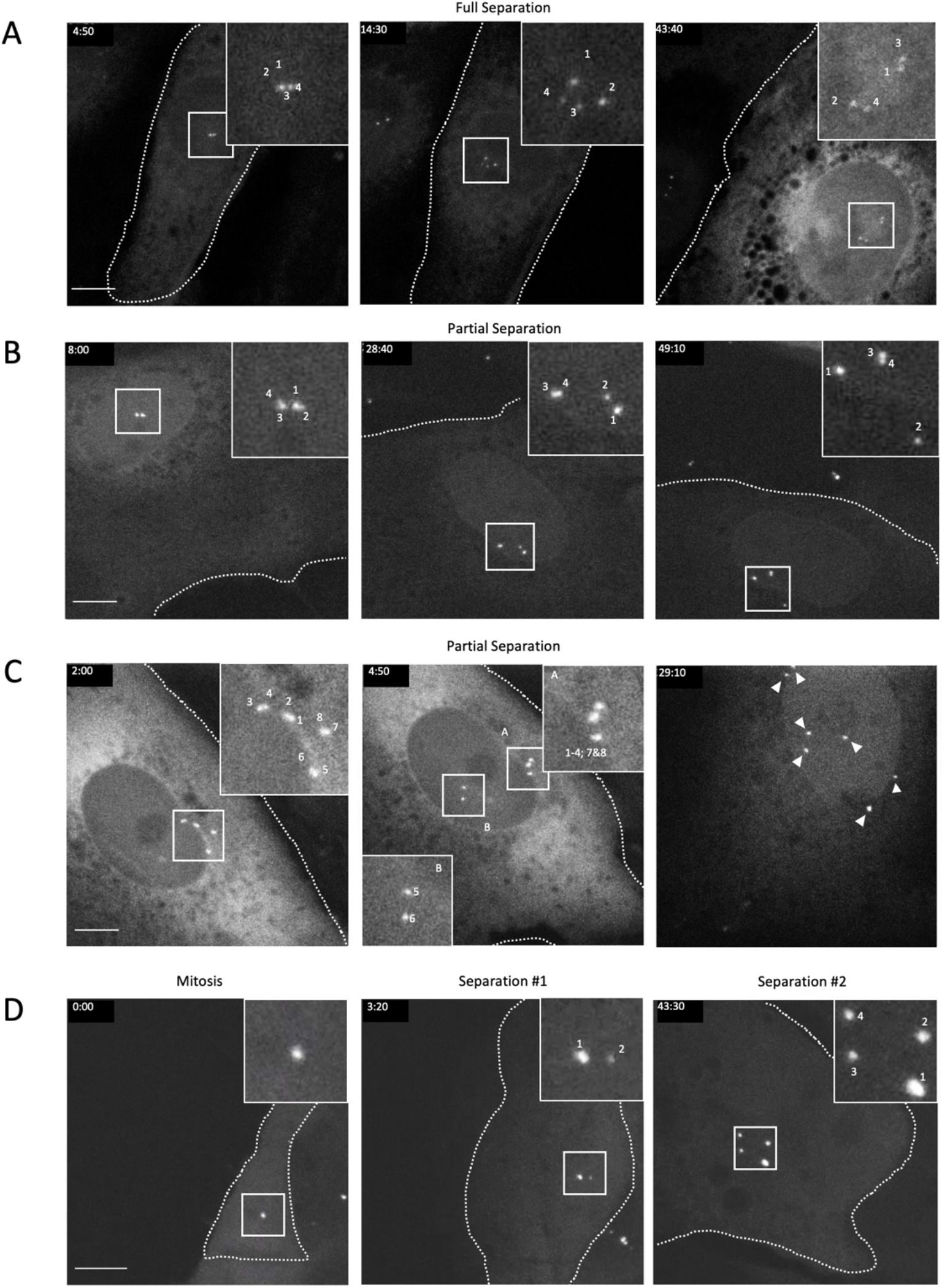
Centriole disengagement in post-mitotic TGCs A-D) Differentiating TGCs, with centrioles marked by eGFP-centrin2, demonstrating disengagement and full or partial separation of duplicated centrioles. Scale bars = 10 µm. A. Still frames from Movie S1 demonstrating full separation of both centriole pairs. Panel 1 shows engaged configuration of centrioles that persists for ∼14 hours before disengaging and separating both pairs of centrioles in panel 2 “Full Separation”. Panel 3 shows the same four centrioles nearly 30 hours later while the cell has dramatically enlarged both its nucleus and cytoplasm. Insets are shown at 3x magnification B. Still frames from Movie S2 demonstrating partial separation of a single centriole pair. Panel 2, “Partial Separation” demonstrates a single pair of centrioles (“3-4”) undergoing disengagement and separation while the other pair (“1-2”) remains tightly associated. Panel 3 shows the same cell with centrioles 3 and 4 remaining separated and centrioles 1-2 remaining associated hours later. Insets are shown at 3x magnification C. Still frames from Movie S3 demonstrating partial centriole separation for a pair of engaged centrioles in a cell with supernumerary centrioles. Panel 2, “Partial Separation” demonstrates a single pair of centrioles (“5-6”) undergoing disengagement and separation while the other pairs (“1-4; 7&8”) remain associated. Panel 3 shows the same cell one day later with most centrioles separated throughout the cell (arrowheads). Insets are shown at 2x magnification D. Still frames from Movie S4 demonstrating duplication, disengagement and separation after a mitotic event. Panels show a TSC undergoing mitosis (first panel, “Mitosis”), disengaging and separating centrioles (“Separation #1”), duplicating, disengaging, and separating centrioles again (“Separation #2”) without passing through a second mitosis. Insets are shown at 3x magnification.

In 95 time-lapse sequences, the centrioles in most cells behaved as expected for the canonical centriole duplication cycle in mitotically dividing cells. However, in a fraction of cells, which we assumed to be TGCs based on their position at the edge of a colony and increasingly large area and nuclei, we observed non-canonical centriole events relevant to the two major TGC centrosome phenotypes of centriole disengagement and amplification.

We found that in some cells (n=7 examples) paired centrin foci separated (distance ≥ 2 µm) without an intervening mitosis (Figure 3A, B and D), consistent with the enrichment for single-centriole centrosomes observed in fixed cells (Figure 2B-D). In some cases the separation of centrin foci occurred relatively close in time (∼1 h) (Figure 3A), whereas in others, separation occurred further apart in time (>20 h) (Figure 3B,D) despite the centriole pairs being in a common cytoplasm.

Less frequently (n=3 examples), we found supernumerary centrin foci that appeared as pairs, consistent with the normal pattern of centriole duplication in which each parental centriole begets a single new centriole. In the example shown in Figure 3C, the cell begins with four pairs but rapidly separates a single pair (Figure 3C, numbers 5-6 and Video S3, 3:50) while the rest remain engaged. Approximately 20 h later, another pair separates (Video S3, 24:20). Thus, non-mitotic disengagement and separation can occur in differentiating TGCs with both normal and amplified centriole numbers.

We note that in the example shown in Figure 3D (Supplementary Video 4), we were able to image the entire centriole cycle as a cell transitioned from mitotic cycling to the early stages of TGC differentiation, which culminated in the separation of two pairs of centrioles approximately 40 h after the last mitosis. These results show that differentiating TGCs duplicate and disengage their centrioles without passing through mitosis, and that cells with amplified centrioles are able to disengage their centrioles in the absence of mitosis. Interestingly, separation events of engaged pairs can be tens of hours apart, indicating a single biochemical reaction does not determine separation of all engaged pairs in a shared cytoplasm.

### Amplified centrioles in TGCs acquire microtubule nucleation competence

Given that about half of differentiating TGCs created new centrioles during the differentiation, and that newly formed centrioles typically mature in conjunction with the cell cycle, we sought to characterize the composition and function of centrioles formed within the context of the endocycle. We first used expansion microscopy to visualize centrioles at higher resolution. Cells were stained with antibodies against acetylated ɑ-tubulin, to mark centriolar microtubules, and ***γ***-tubulin, to mark pericentriolar material and centriole lumen. (Gambarotto et al., 2021; Wassie et al., 2019). We found that the centriolar structure in TGCs was similar to previous descriptions of centrioles in human cells imaged by expansion microscopy (Gambarotto et al., 2019). Acetylated tubulin labeling showed cylindrical structures with the dimensions of normal centrioles, some with orthogonally positioned procentrioles, identified from their shorter length (Figure 4A). All centrioles had associated ***γ***-tubulin, both in pericentriolar distribution and in the lumen, as expected for centrioles that have matured to become centrosomes and functional MTOCs (Schweizer et al., 2021).

**Figure 4:**
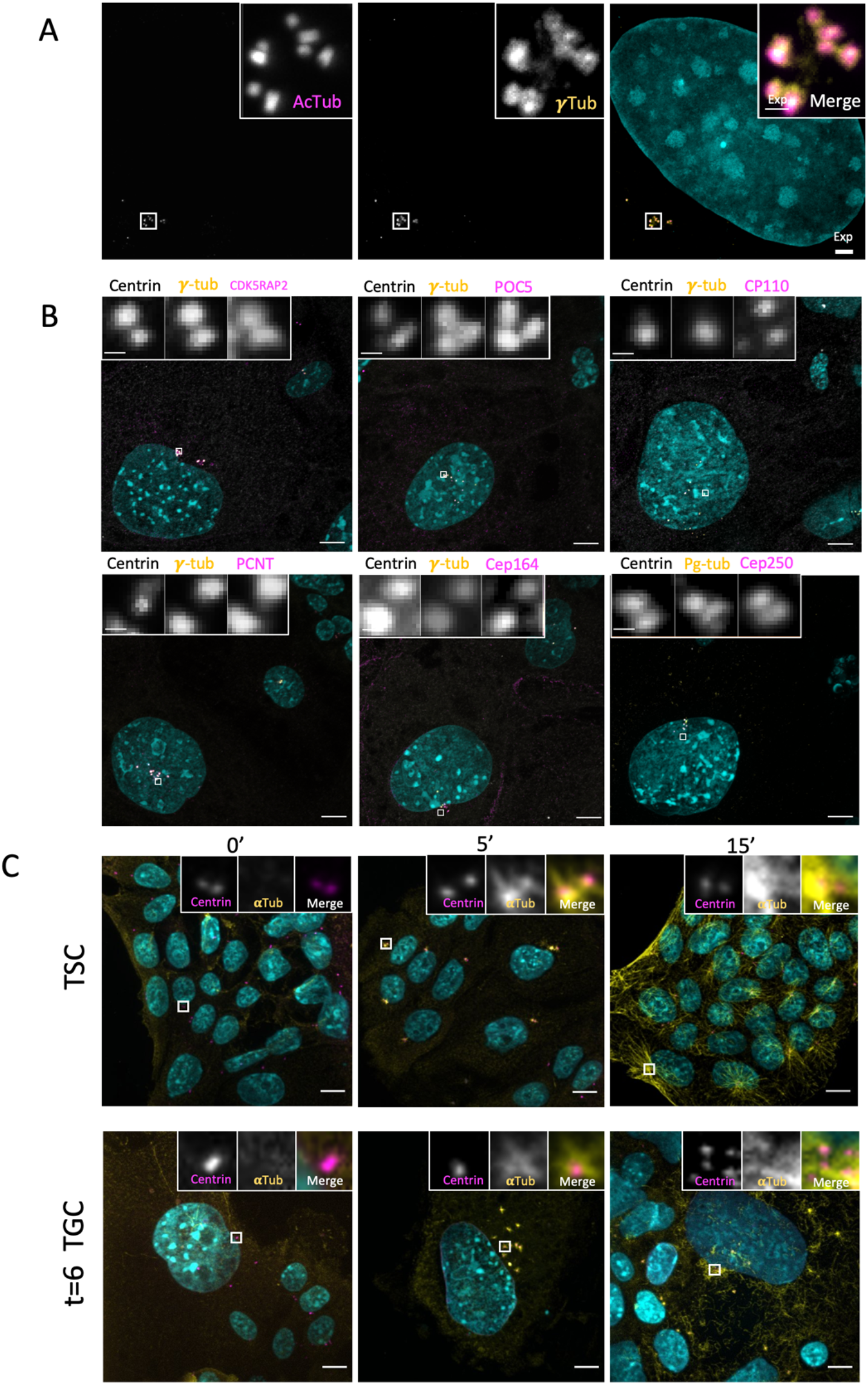
Amplified centrioles in TGCs acquire microtubule nucleation competence A. Expansion microscopy images of TGCs demonstrating supernumerary centriole content with expected morphology. Expanded TGCs were stained with antibodies to mark the centrioles (acetylated tubulin) and PCM (***γ***-tubulin) and were counterstained with DAPI, to mark nuclei. Scale bars = 10 µm, inset = 2 µm (expansion). B. Immunofluorescence images showing localization of various centrosome proteins in TGCs with amplified centrosomes. Cells were fixed and labeled with antibodies to label structural, PCM and appendage proteins as indicated above each panel. Scale bar = 10 µm, inset = 1 µm C. Immunofluorescence images showing TSCs (top) and TGCs (bottom) in a microtubule regrowth experiment. First panel shows cells just prior to washout (0 Min). Second and third panels show cells shortly after washout (5 min and 15 min). Cells were fixed and labeled with antibodies to mark microtubules (α-tubulin, yellow), centrioles (centrin, magenta) and nuclei (DAPI, cyan). A-C Scale bars = 10 µm, insets in C are shown at 7x magnification.

The life cycle of centrioles in mitotically dividing cells is characterized by changes to centriole structure that occur over the course of more than one cell cycle and some of which require passage through mitosis (Izquierdo et al., 2014; Kong et al., 2014; Tsou & Stearns, 2006). We examined centrioles in TGCs for markers indicative of those changes to determine whether they also occur in the TGC endocycles. TGCs at t=8d were examined for tubulin modifications (Figure 4A, acetylated ɑ-tubulin) (polyglutamylated tubulin) proteins of the distal lumen (POC5, centrin), the proximal end (CNAP1) the centriole cap (CP110), pericentriolar material (***γ***-tubulin, Pericentrin, CDK5RAP2) and distal appendages (CEP164) (Figure 4B). For all proteins examined, the centrioles in TGCs had the expected localization of the proteins considering the stage of duplication (i.e. single centrioles vs. centrioles with procentrioles). The one exception was CEP164, which was not present on every centriole but could be readily found on more than one centriole per cell.

Next, we tested whether amplified centrioles were competent to serve as MTOCs, using nocodazole for microtubule depolymerization and washout to visualize newly-nucleated microtubules during regrowth. TSCs and t=6d TGCs were treated with 10 µg/ml nocodazole for 1h to depolymerize microtubules and assessed for regrowth at timepoints following washout (ɑ-tubulin) (Figure 4C), as well as the location of centrioles (centrin). Microtubule asters began to grow from centriole foci in TSCs within 5 minutes, with no more than two asters per cell. In TGCs, microtubule asters (Figure 4C) formed from each centriole focus, including in TGCs with amplified centrioles, suggesting that each centriole focus, even those consisting of single centrioles, is capable of serving as a functional MTOC. Thus, centrioles formed during the endocycle in TGCs undergo the centriole-to-centrosome conversion, assemble PCM, acquire maturity markers, and serve as functional MTOCs.

### PLK4 activity is required for centriole amplification, but not for reduplication of DNA during the TGC endocycle

Since centriole and centrosome number increases with differentiation, we sought to determine if centriole formation was necessary for endocycles to proceed in differentiating TGCs. Although it is not currently possible to acutely prevent centriole separation and centrosome formation, we could prevent the formation of additional centrioles by treatment with Centrinone-B, an inhibitor of PLK4, the master regulator of centriole biogenesis (Holland et al., 2012; Sillibourne & Bornens, 2010; Wong et al., 2015). TSCs were treated with Centrinone-B at the beginning of differentiation and grown for 6 days (Figure 5A). Using centrin as a marker of centrioles, we determined the number of centrioles per cell relative to EdU status (Figure 5B-E). Centrinone-B treatment reduced the number of centrioles per cell, including greatly reducing instances of cells with four or more centrioles, suggesting that PLK4 inhibition was effective (Figure 5B,C). This suggests that centriole increase in TGCs is PLK4-dependent, just like in mitotic cells. To assess whether TGCs amplified their DNA despite the inhibition of centriole formation, newly synthesized DNA was labeled by incubation with the analog 5-ethynyl-2’-deoxyuridine (EdU) 24 hours prior to the endpoint of the experiment (Figure 5A). Centrinone-B treatment did not significantly alter nuclear size or the fraction of EdU-positive cells (Figure 5D,E). We note that previous work suggested that PLK4 is required for TGC differentiation via phosphorylation of the transcriptional regulator HAND1 (Hemberger et al., 2004; Martindill et al., 2007). Given a more recent understanding of the essential role of PLK4 in centriole duplication and the consequences of centriole loss, we suggest that the earlier results might instead be due to centriole loss. These results suggest that PLK4 activity is necessary for increased centriole number but is dispensable for progression through the endocycle during TGC differentiation.

**Figure 5:**
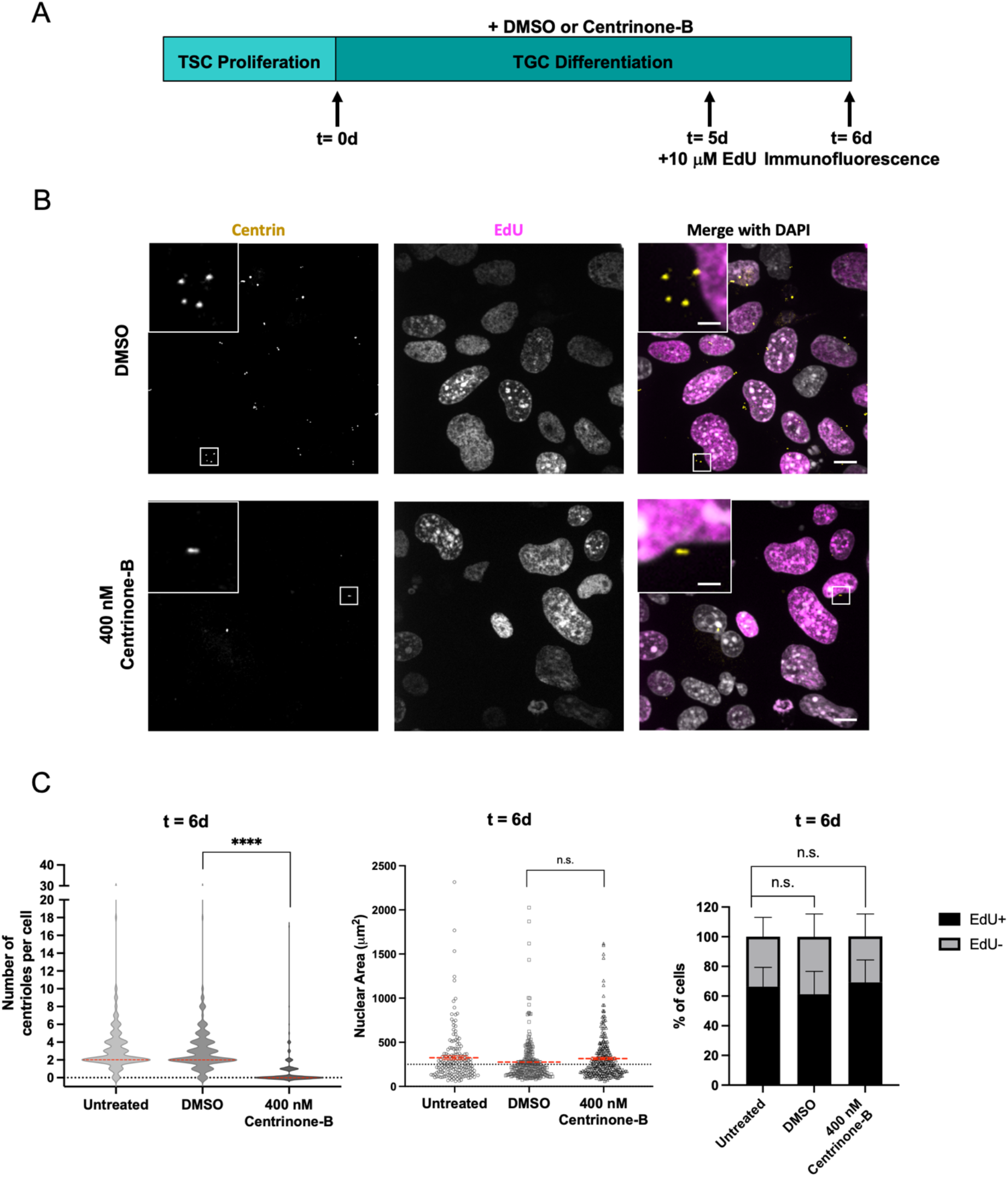
Increase in TGC centriole number during differentiation is dependent on PLK4 activity A. Schematic of experimental design. TSCs were seeded 24 hours prior to differentiation, as indicated by TSC proliferation. TGC differentiation was conducted in the presence of Centrinone-B, or an equivalent volume of DMSO as a control for up to six days. An untreated control was also included. To label nuclei that are actively replicating DNA in S-phase, TGCs were incubated with 10 µM 5-ethynyl-2’-deoxyuridine (EdU) at 5d for 24 hours. B. Immunofluorescence of representative images of TGCs at t=6d. Centrioles were labeled with centrin (yellow) antibody and nuclei that entered S-phase were labeled for EdU incorporation using Click-it® Chemistry (magenta). Scale bars 10 µm inset = 2 µm C. Violin plot of centriole number in cells for t=6d during drug treatments and control conditions. Solid red lines indicate median. D. Quantification of nuclear area for t=6d during drug treatments and control conditions. Red horizontal lines represent the mean. Each dot represents a single cell mean (in red) E. Quantification of the EdU status for populations of cells at t=6d during drug treatments and control conditions. Results shown are for three independent experiments. At least 50 cells quantified per experiment for untreated, DMSO, and Centrinone-B treatments, error bars are the SEM. ****p-value ≤0.0001, n.s., not significant.

## Discussion

Centriole formation usually occurs in the context of mitotically dividing cells, where it is coupled to DNA replication and cell cycle progression. It can be uncoupled from these events in specialized differentiated cells that make many centrioles under the control of a transcriptional program (Kyrousi et al., 2015; Spassky & Meunier, 2017; Vanderlaan et al., 1983) without replicating DNA. Here we have examined endocycling cells of the trophoblast lineage to determine whether centriole formation is coupled or uncoupled to the DNA endocycles. We find that TGCs *in situ* have amplified centrioles and that this can be recapitulated in an *in vitro* differentiation system. However, centriole number in these cells does not increase exponentially with DNA reduplication, as would be expected for tight coupling of these two processes. Rather, centrosome number increases to a greater extent than centriole number in TGCs because centrioles that are duplicated disengage, separate and become centrosomes, such that each centriole is an independent, functional microtubule organizing center. We consider the implications of these findings for understanding centriole and centrosome number control in division and differentiation.

In Figure 6 we propose a model that integrates our findings. A TSC (shown on left) with two centrosomes (yellow dots) with either two or four centrioles (magenta barrels) either divides and cycles or undergoes differentiation into a TGC (shown on right). In their final state, TGCs have disengaged and separated centrioles that have acquired centrosome competence to reach a centriole to centrosome ratio of 1:1 (cell 2 and cell 4). In some cases, the total number of centrioles in a TGC is equal to that in the TSC from which it is derived, after one round of duplication (cell 2; four centrioles), whereas in others further rounds of centriole formation have occurred (cell 4; > four centrioles). Only disengaged centrioles are competent to duplicate, and we observe that disengagement and separation does not occur synchronously, resulting in centriole/centrosome numbers that are not a simple function of 2^n^. Additionally, our data cannot rule out the possibility that centrioles may also arise by some mechanism other than canonical duplication or that procentrioles formed by canonical duplication are not occasionally degraded. Many TGCs appeared to have odd numbers of centrioles and centrosomes rather than the even number expected from this model, and this may have resulted from such processes, but could also have been the result of inherent difficulties of imaging centrioles. In any case, TGCs bearing extra centrioles also ultimately resolve to a 1:1 ratio of centrioles to centrosomes.

**Figure 6:**
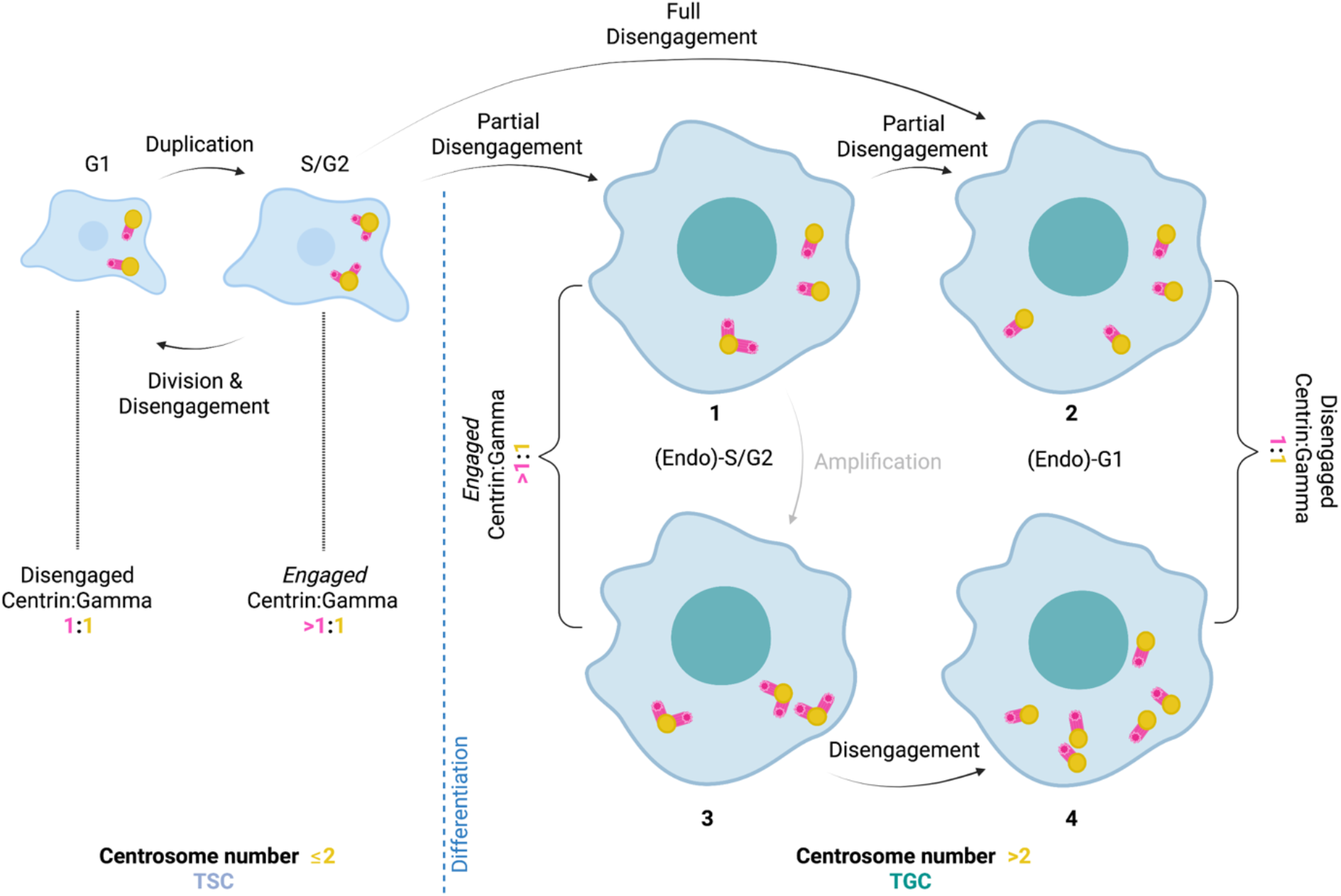
Model of Centriole and Centrosome Amplification in Endocycling Murine TGCs Cartoon diagram showing (left) a TSC with two centrosomes (yellow dots) with either two or four centrioles (magenta barrels) and ratios of ***γ***-tubulin (to identify centrosomes) and centrin (to identify centrioles) given below. (Right) TGCs during differentiation with a variable number of centrosomes (yellow dots) and centrioles (magenta barrels). Most TGCs disengage their paired centrioles to reach a centrin to ***γ***-tubulin ratio of 1:1. Some TGCs undergo additional centriole/centrosome amplification that also ultimately resolves to a 1:1 ratio of centrin to ***γ***-tubulin at the end of the endocycle. Graphic was created with Biorender.com.

In cycling cells, disengagement of the procentriole from the mother centriole is a rate-limiting step for forming new centrioles. Indeed, laser ablation of a procentriole results in a new centriole forming from the same mother centriole (Lončarek et al., 2010). In contrast, we found that TGCs could exist for prolonged periods (days) with fully disengaged centrioles and no further centriole formation. One explanation would be that the transcriptional program of the TGC endocycle either does not express the genes for centriole duplication or expresses an inhibitor of the process. We note that the canonical centriole duplication proteins SASS6, PLK4 and STIL are all significantly downregulated in t=4d TGCs as determined by RNAseq (Ullah et al., 2020). How then would the relatively small fraction of TGCs with greater numbers of centrioles arise? It could be that these cases are due to the protein remaining after initiation of TGC differentiation or might be due to stochastic variation in establishment of the TGC transcriptional program or incomplete repression of expression of centriole duplication genes could be sufficient to allow for duplication in some cells (Raj & Van Oudenaarden, 2008). This is consistent with the observation that most TGCs that have more than four centrioles have only five or six centrioles, rather than, for example, eight, as would be expected for another complete round of duplication.

There are several differentiated cell types in mammals that break the “once and only once” pattern of centriole duplication observed for most cycling cells. Examples include multiciliated cells of the airway, chorid plexus, oviduct, and olfactory sensory neurons (Ching & Stearns, 2020; Klos Dehring et al., 2013; Narita & Takeda, 2015; Spassky & Meunier, 2017). In most of these cell types the amplified centrioles form cilia, and the function of these cells is dependent on having multiple cilia (Ching & Stearns, 2020; Klos Dehring et al., 2013; Narita & Takeda, 2015). TGCs do not form even a single primary cilium, so presumably the observed amplification is not related to cilium functions. A unique feature of TGC centriole formation is that it mostly occurs by the usual process of initiating one procentriole on the side of a mother centriole, but that the centriole pairs all disengage, separate and become centrosomes, maximizing the number of centrosomes. Might this greater number of centrosomes be related to the properties of TGCs? Recent reports show that the presence of extra centrosomes in mammalian cells causes a range of phenotypes, including increased invasiveness (Godinho & Pellman, 2014). Thus, a possible explanation for the function of the extra centrosomes in TGCs is that they aid in promoting the invasive phenotype of TGCs, which is critical to their ability to migrate into the placenta during pregnancy. Another possibility is that the extra centrosomes in TGCs are not necessary for TGC function, but instead are simply a byproduct of this form of endocycling. Whether extra centrosomes are required for the TGC function will require testing *in vivo*, using cell-type-specific genetic manipulation to block centriole formation.

The centriole and centrosome events that we have observed in TGC endocycles bear a remarkable resemblance to those described in some mammalian cell types under prolonged S-phase arrest. Balczon, et al. (1995) showed that arrest of CHO cells in S-phase with hydroxyurea resulted in amplification of centrioles over several days. Subsequent work on this S-phase arrest amplification has shown that even among transformed cell lines this is not a universal property and instead seems to be associated with PLK1 activity during the prolonged arrest (Lončarek et al., 2010). Centrioles duplicated under these conditions often disengage and can become competent for further duplication (Balczon et al., 1995; Lončarek et al., 2010). Disengagement in a typical cell cycle is likely a multistep process, initiated in G2 and completed in late mitosis (Lončarek et al., 2010; Tsou & Stearns, 2006). However, in both the TGC endocyle and prolonged S-phase, arrest can occur without transiting mitosis. Also, in both cases disengagement can occur asynchronously with engaged centriole pairs separating hours apart (Lončarek et al., 2008). It remains to be determined what centriole-intrinsic factors in TGCs might be responsible for this asynchronous behavior in a common cytoplasm, but we note that there are many examples of localized phenomena in shared cytoplasm, such as the asynchronous division of nuclei in the filamentous phase of the fungus *Ashbya* (Gladfelter et al., 2006).

In summary, we find that murine TGCs differentiating *in vitro* undergo endocycles that increase DNA content without substantially increasing centriole number in most cells beyond the normal S/G2 number of four. However, the centrioles that do exist are disengaged and separated and become centrosomes, such that most TGCs have four or more functional centrosomes. It will be of interest to determine whether the many other examples of genome amplification in differentiated cells are similar or different in regard to centrosome properties and the function they might provide.

## Acknowledgements

This project was supported by the NIGMS of the National Institutes of Health under award numbers 1R35GM130286 (to T.S.) and T32GM007276 (to G.B and M.S.). We thank Julie Baker’s lab for the gift of TS cells and all members of the Stearns lab for helpful feedback and suggestions.

## Competing Interests

We do not declare any competing interests at this time.

## Author Contributions

Conceptualization and writing: G.B., M.S., L.M., and T.S.; Investigation: G.B., M.S., and L.M.; Supervision, funding acquisition, and project management: T.S.

## Materials and Methods

### Ethics Statement

This study uses samples from mice. All animal procedures in this study were approved by the Stanford University Administrative Panel for Laboratory Animal Care (SUAPLAC protocol 11659) and carried out according to SUAPLAC guidelines.

### Cell lines

Mouse trophoblast stem cells (TSCs) were a gift from Julie C. Baker (Stanford University) and were maintained in DMEM-F12 with 15 mM HEPES (Gibco, Life Technologies, catalog # 11330032), 20% fetal bovine serum (Gemini Bioproducts, catalog #900-208, lot # A00G91I), 2 mM Glutamax (Gibco, Life Technologies, catalog # 35050061), 100 µg/ml penicillin-streptomycin, 1 mM sodium pyruvate (HyClone, GE Healthcare Life Sciences, catalog # SH3023901), 100 µM β-mercaptoethanol, and 1X MEM nonessential amino acids, supplemented with 10 ng/ml Activin A (Peprotech, catalog # 120-14P), 25 ng/ml fibroblast growth factor 4 (FGF4, Peprotech, catalog #100-31), and 1 µg/ml heparin (Sigma-Aldrich), as described in (Chuong et al., 2013; Tanaka et al., 1999).

To differentiate TSCs into parietal trophoblast giant cells (TGCs), TSCs were seeded at 2.5×10^4^ cells per well in 6-well or 12-well plates. The following day, TSCs were differentiated into TGCs by removing the FGF4, Activin A, and heparin from growth media, and replacing with 5 µM retinoic acid. TGCs were maintained for up to 10 days in culture, where media was changed every two days.

### Mouse husbandry and derivation of TSCs

Arl13b-mCherry;eGFP-centrin2 transgenic mice (JAX#027967) were obtained from JAX laboratories, which were generated by (Bangs et al., 2015) on FVB and C3H mixed background. All procedures involving animals were approved by the Institutional Animal Care and Use Committee of Stanford University School of Medicine in accordance with established guidelines for animal care. Breeding pairs were mated to establish timed pregnancies. Copulation was determined by the presence of a vaginal plug the morning after mating, and embryonic day 0.5 (e0.5) was defined as noon of that day. Embryos ranging from e3.5d to e9.5d, as well as adult males and females for breeding between 8 weeks and 9 months were used for this study.

Arl13b-mCherry;eGFP-centrin2 TSCs were derived from mouse blastocysts isolated from pregnant mice at e3.5d as described by (Kidder, 2014; Tanaka, 2006). Initially, blastocysts were seeded onto a feeder layer of irradiated mouse embryonic fibroblasts (iMEFs) after isolation, then observed for an outgrowth of cells. Arl13b-mCherry;eGFP-centrin2 TSCs were maintained in the same growth conditions as TSCs stated above. Fetal bovine serum used for TSC derivation and maintenance was purchased from HyClone (GE Healthcare Life Sciences, catalog # SH30070.02, lot# AC1024054S).

### Immunofluorescence and immunohistochemistry

TSCs and TGCs were grown on poly-L-lysine–coated #1.5 glass coverslips (Electron Microscopy Sciences, Hatfield, PA) for confocal microscopy. TSCs and TGCs were washed with phosphate buffered saline (PBS), fixed in -20°C methanol for 10 minutes, washed three times with PBS, then blocked in 3% bovine serum albumin (BSA), 0.1% Triton X-100, and 0.02% sodium azide in PBS (PBS-BT) for 30 minutes to 1 hour at room temperature. Samples were then incubated with primary antibodies overnight at 4°C. The following day, samples were washed three times in PBS-BT, then incubated with Alexa Fluor dye-conjugated secondary antibodies (Invitrogen) diluted 1:1,000 in PBS-BT for 1 hour at room temperature. When applicable, appropriate isotype-specific secondary antibodies were used to distinguish different monoclonal mouse antibodies. Samples were then stained with 5 µg/ml DAPI (4′,6-diamidino-2-phenylindole) for two minutes to visualize nuclei and mounted in Mowiol mounting medium (Polysciences) in glycerol containing 2.5% 1,4-diazabicyclo-(2,2,2)-octane (DABCO, Sigma-Aldrich) antifade.

The conceptus was isolated from a pregnant mouse at e9.5d and fixed in 4% paraformaldehyde at 4°C overnight. The conceptus was then washed three times in PBS, incubated in 30% sucrose at 4°C overnight for cryoprotection, embedded in OCT compound and frozen at -80°C. The conceptus was then cryosectioned at 10 µm thick sections. For immunohistochemistry, tissue was rehydrated with 1% normal goat serum in PBS for 30 minutes at room temperature. Nuclei were stained using DAPI as indicated in the immunofluorescence protocol above.

### Antibodies

Primary antibodies used were mouse monoclonal anti-***γ***-tubulin (Sigma-Aldrich, clone GTU88, IgG1; used at 1:5,000 dilution in PBS-BT), mouse monoclonal anti-centrin3 (Novus Biologicals, clone 93E6, IgG2b; 1:2,000), mouse monoclonal anti-α-tubulin (Abcam, clone DM1α, IgG1; 1:4,000, St. Louis, MO), mouse monoclonal anti-centrin (Sigma-Aldrich, clone 20H5, IgG2a; 1:200), mouse monoclonal anti-acetylated tubulin (Sigma-Aldrich, clone 6-11B-1, 1:1000) rabbit polyclonal anti-POC5 (A303-341A-T, Bethyl Laboratories, 1:500), rabbit polyclonal anti-CP110 (12780-1-AP, Proteintech, 1:200), rabbit polyclonal anti-CNAP1/Cep250 (Proteintech, 14498-1-AP, 1:500), rabbit polyclonal anti-Pericentrin (Abcam, ab4448, 1:500), rabbit polyclonal anti-CDK5RAP2 (Sigma-Aldrich, 06-1398, 1:1000), and rabbit polyclonal anti-Cep164 (purified from serum of immunized animals as described in (Lau et al., 2012), 1:1,000).

### Drug treatments, EdU labeling and Centrinone-B treatment

For inhibition of centriole duplication experiments, 400 nM Centrinone-B, or the equivalent volume of DMSO as a vehicle control were added to TSCs upon inducing differentiation. TGCs were maintained in the presence of these treatments throughout the duration of the time course. 10 µM EdU (5-ethynyl-2’-deoxyuridine) was added 24 hours before the termination of a time course at day 5 post-differentiation to examine if DNA replication occurred during drug treatment. Staining for EdU incorporation was conducted using Click-iT® EdU Alexa Fluor® 594 Imaging Kit (Life Technologies, catalog #C10339), followed by immunofluorescence as mentioned above.

### Microtubule regrowth assay

TSCs were seeded onto coverslips at 5×10^4^ cells in 3.5 cm dishes and differentiated the following day. To depolymerize microtubules, cells were treated with 10 µg/ml nocodazole for 1 hour at 37°C. Cells were then washed with ice cold PBS on ice. Microtubule regrowth was then initiated when coverslips were incubated with medium warmed at 37°C. To stop the assay, cells were fixed with 100% methanol for 10 minutes at -20°C at the respective time points before proceeding with immunofluorescence as described above.

### Confocal Microscopy and live cell imaging

Confocal microscopy images were acquired as Z-stacks collected at 0.5-μm intervals across a 15 – 30 μm range for fixed cells on a Zeiss Axio Observer microscope (Carl Zeiss) with a confocal spinning-disk head (Yokogawa Electric Corporation, Tokyo, Japan), PlanApoChromat 63×/1.4 NA objective, and a Cascade II:512 electron-multiplying (EM) CCD camera (Photometrics, Tucson, AZ) or PRIME: BSI backside illuminated CMOS camera run with µ-Manager software (Edelstein et al., 2014) or SlideBook 6 software (3i, Denver, CO).

Confocal microscopy images for fixed cells were also acquired using a Leica SP8 scanning confocal microscope with a 63 × (1.4 N.A.) objective at the same Z-stack parameters listed previously. All images were processed using Fiji (National Institutes of Health, Bethesda, MD) and/or SlideBook (3i, Denver, CO).

The medium was changed 1 hour prior to imaging to phenol-free DMEM-F12 with 15 mM HEPES supplemented with 20% fetal bovine serum, 100 µg/ml penicillin-streptomycin, 1 mM sodium pyruvate, 100 µM β-mercaptoethanol, 1X MEM nonessential amino acids, and 5 µM retinoic acid. During image acquisition, cells were incubated at 37°C under 5% CO_2_. Images were acquired as 0.5 µm Z-stacks collected every 5 or 10 min for up to 72 hours using a Zeiss Axio Observer microscope (Carl Zeiss) with a confocal spinning-disk head (Yokogawa Electric Corporation, Tokyo, Japan), PRIME: BSI backside illuminated CMOS camera run with µ-Manager software (Edelstein et al., 2014) or SlideBook 6 software (3i, Denver, CO).

### Expansion Microscopy

Expansion microscopy was performed according to the previously described U-ExM protocol (Gambarotto et al., 2019). In brief, TGC cultures on coverslips were fixed in ice cold methanol and incubated for 10 minutes at –20°C for 10 min, then washed with 1× PBS. PBS was removed and cells were incubated in monomer fixative solution (0.7% formaldehyde and 1% w/v acrylamide in water) for 4-5 hr at 37°C. Coverslips were then inverted onto droplets of cold gelation solution (19% w/v sodium acrylate, 10% w/v acrylamide, 0.1% BIS, 0.5% TEMED, 0.5% ammonium persulfate in PBS) and allowed to set for 5 min on ice. Coverslips were then transferred to 37°C for 1 hr. Coverslips with gels were incubated in denaturation buffer (200 mM SDS, 200 mM NaCl, and 50 mM Tris in water) for 15 minutes at RT to allow the gel to detach from the cover slip and then the gel was transferred to a 1.5ml Eppendorf with denaturation buffer for 45 min at 95°C. Gels were removed from the Eppendorf and washed in water twice for five minutes and then expanded in water overnight. Gels were then incubated in PBS-BT for 30 min prior to any staining. Gels were incubated in primary antibody solution in PBS-BT on a nutator for 3-4 hr to overnight. Next, gels were washed three times in PBS for 10–30 min per wash and then incubated in a solution of secondary antibodies conjugated to Alexa Fluors plus DAPI, diluted 1:1000, overnight at 4°C. Gels were washed in PBS for at least 30 min, then in water three times for 10 min per wash. Gels were then allowed to fully expand in water for at least an hour before mounting in a glass-bottom imaging dish and imaging by spinning disk confocal microscopy.

### Quantification of nuclear area and statistical analyses

Nuclear area and integrated DAPI measurements of mononucleate TGCs were quantified using either maximum intensity or sum projections of stacks acquired by confocal microscopy. Nuclei watershed separation was generated in Fiji on DAPI-stained nuclei as stated on imagej.net (https://imagej.net/Nuclei_Watershed_Separation). Briefly, a Gaussian Blur 3D filter was applied with a sigma value of 3.0 pixels in x, y, and z. Automated threshold plus watershedding was applied to separate individual nuclei. Nuclear area particle analysis parameters were the following: size-50 to 2,000 µm^2^, as reported in (Simmons et al., 2007), and circularity-0.0 to 1.0. If necessary, the upper limit of nuclear area was increased to 5,000 µm^2^. If a single nucleus was segmented into multiple pieces, the area of each segment was added to quantify the nuclear area of the entire nucleus. Nuclei on the edges of the field of view, in addition to binucleate and multinucleate syncytiotrophoblasts, were excluded from analysis (Fig S1A).

Statistical analyses and graphs were generated using GraphPad Prism 9 Software. Pairwise comparisons were made using a one-way ANOVA test. For multiple comparisons, a Dunnett’s test was applied to correct for multiple hypothesis testing. For all analyses, a p-value less than 0.05 was considered significant. Error bars on graphs represent standard error of the mean (SEM), and horizontal lines represent the mean.

### Quantitative real time PCR

RNA was extracted using phenol-chloroform extraction (TriZOL® Reagent, Invitrogen, catalog # 15596-026) from TGCs initially seeded at 2.5×10^4^cells per well in 6-well plates at various time points throughout the differentiation time course indicated. cDNA was prepared from total RNA using Maxima First Strand cDNA Synthesis Kit (Life Technologies, catalog #K1641). qPCR was performed in triplicate with Luna® Universal qPCR SYBR Green Master Mix (New England BioLabs, catalog #M3003L), where 25 ng cDNA was loaded per well. Gene expression was evaluated using the ΔΔCt method (Schmittgen and Livak, 2008). Gapdh levels were used to normalize target gene expression values. For TGC differentiation time courses, gene expression levels were compared to TGCs collected at the beginning of differentiation (t=0d). For validation of derived TSCs from Arl13b-mCherry;eGFP-centrin2 mice, gene expression levels were normalized to TSCs not derived in this paper.

Primers were validated as intron-spanning using PrimerBLAST (NCBI). Primer sequences are indicated below. N/A: not applicable, designed by author using PrimerBLAST-

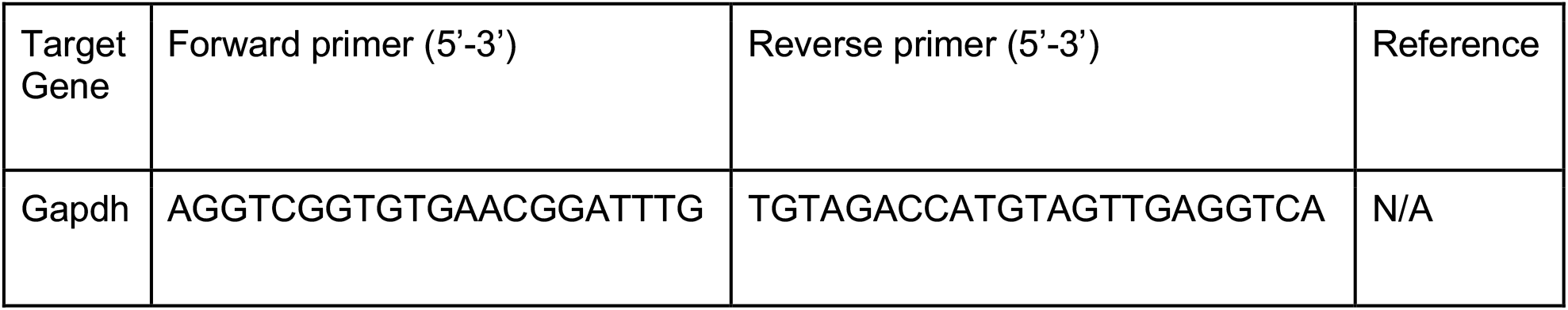
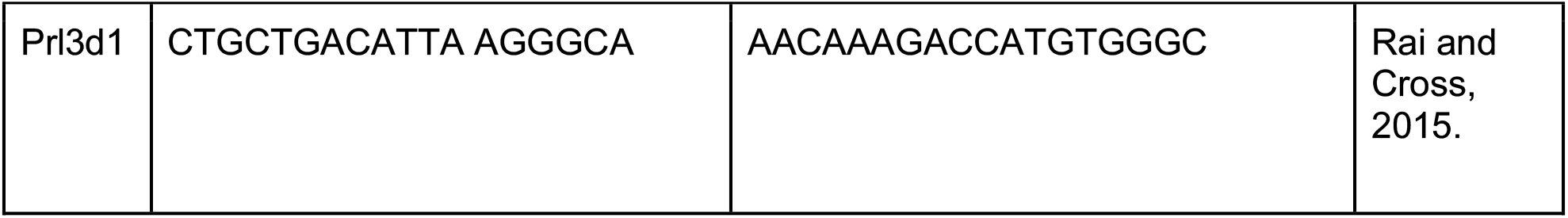

---

**Supplementary Figure 1:**
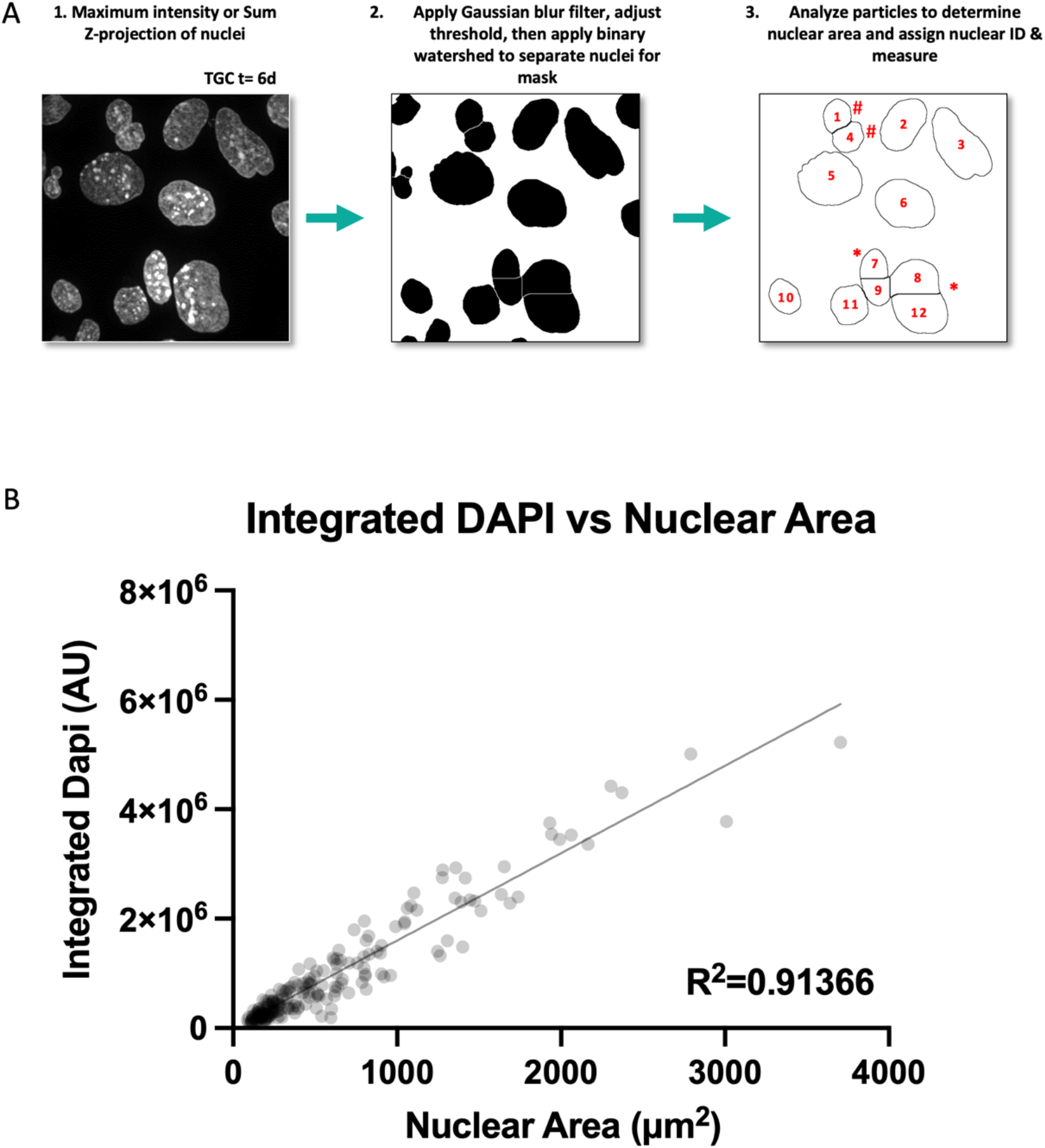
Measurement of Integrated DAPI Signal and Nuclear Area in TGCs A. Schematic of nuclear area quantification from ImageJ. An asterisk (*) indicates nuclear areas that were combined due to an improper division of nuclei relative to the DAPI staining. An octothorpe (#) indicates properly split nuclear areas that are in the same cell that were excluded from analysis B. Plot of the integrated DAPI intensity vs the nuclear area for differentiating TGCs. Each dot on the scatterplot is shaded at 10% opacity and represents a single cell.

**Supplementary Figure 2:**
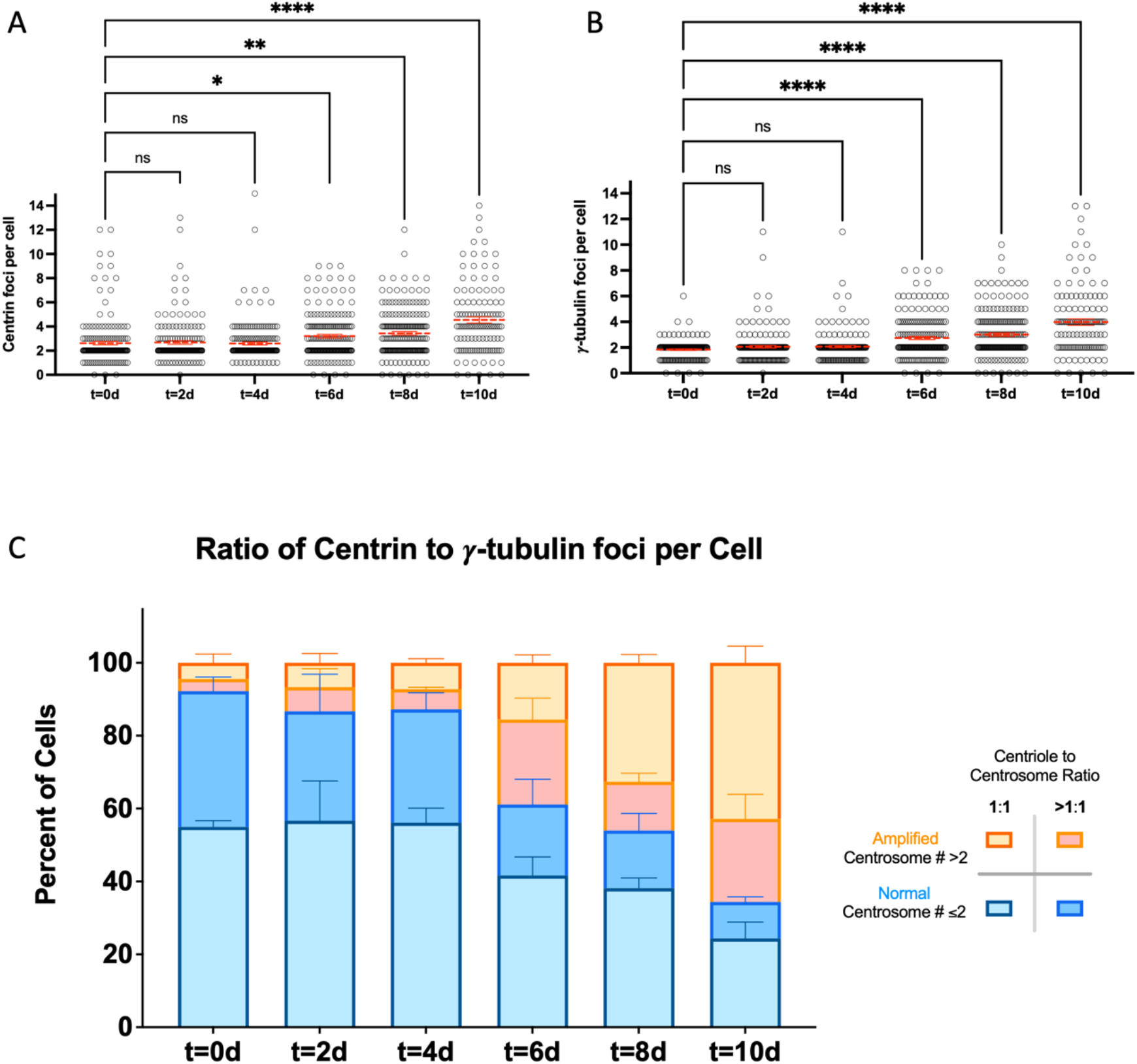
Centrioles and Centrosomes during TGC differentiation A. Quantification of centriole number throughout TGC differentiation as measured by centrin immunofluorescence from same raw data as Figure 2C. Solid red line indicated mean; error bars represent the standard error of the mean (SEM). B. Quantification of centrosome number from (B) throughout TGC differentiation as marked by ***γ***-tubulin immunofluorescence from the same raw data as Figure 2D. Solid red line indicated mean; error bars represent the standard error of the mean (SEM). C. Ratio of centriole and centrosome number, as identified by foci of centrin and ***γ***-tubulin immunofluorescence, respectively, throughout differentiation from the data in (A) and (B).

**Supplementary Video 1:**
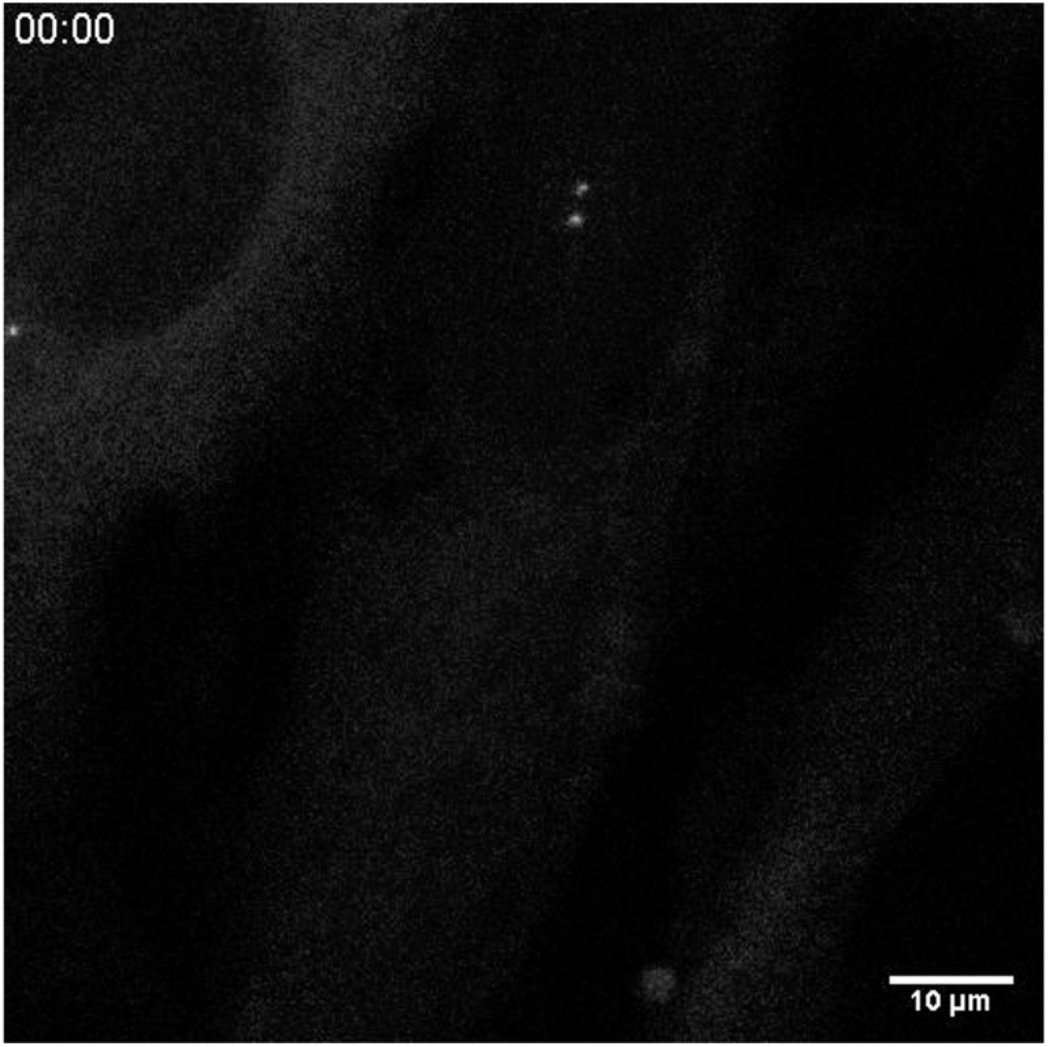
Full centriole disengagement in a differentiating TGC Time lapse movie showing a cell undergoing full centriole separation of both engaged pairs; arrow shows moment of separation. 10 minutes per frame, scale bar is 10μm.

**Supplementary Video 2:**
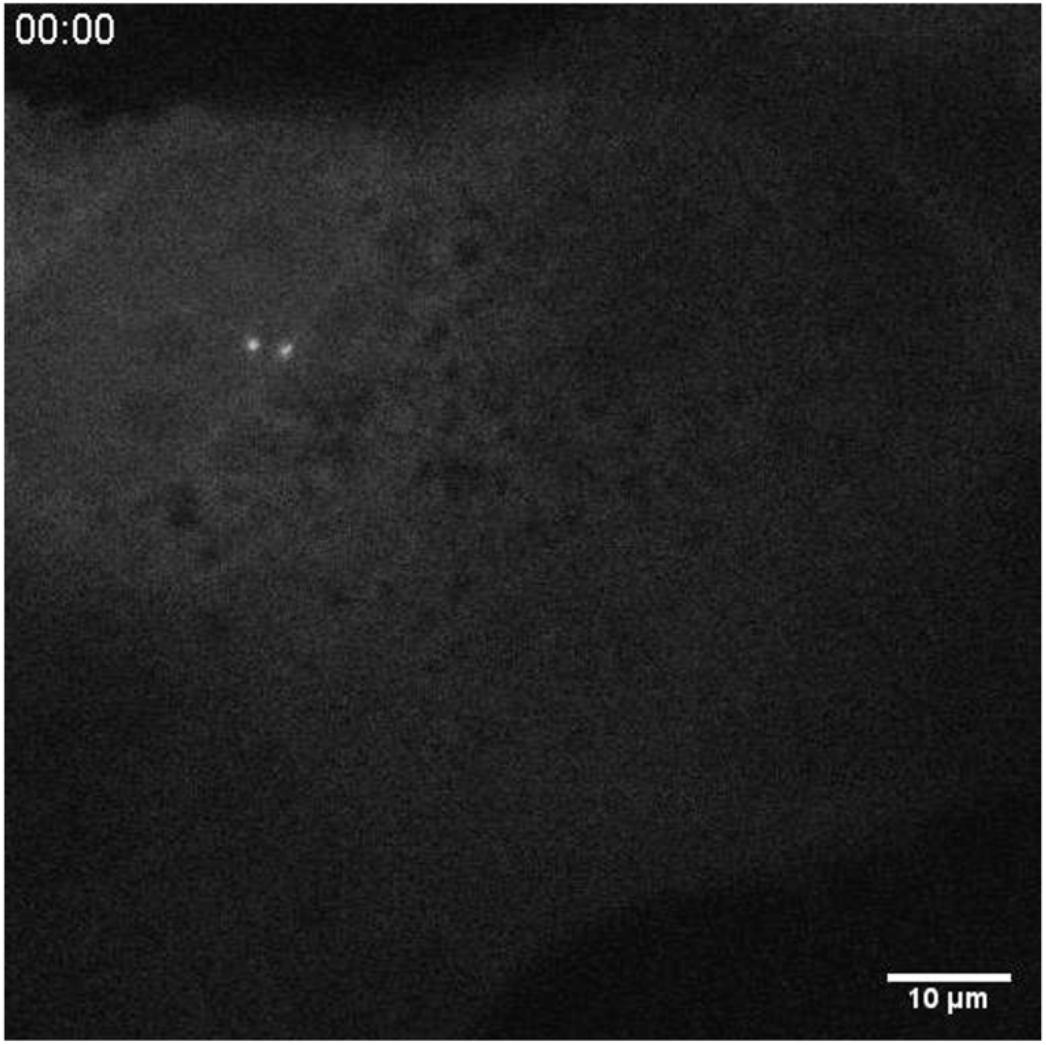
Partial centriole disengagement in a differentiating TGC Time lapse movie showing a cell undergoing partial centriole separation with one of two engaged pairs; arrow shows moment of separation. 10 minutes per frame, scale bar is 10μm.

**Supplementary Video 3:**
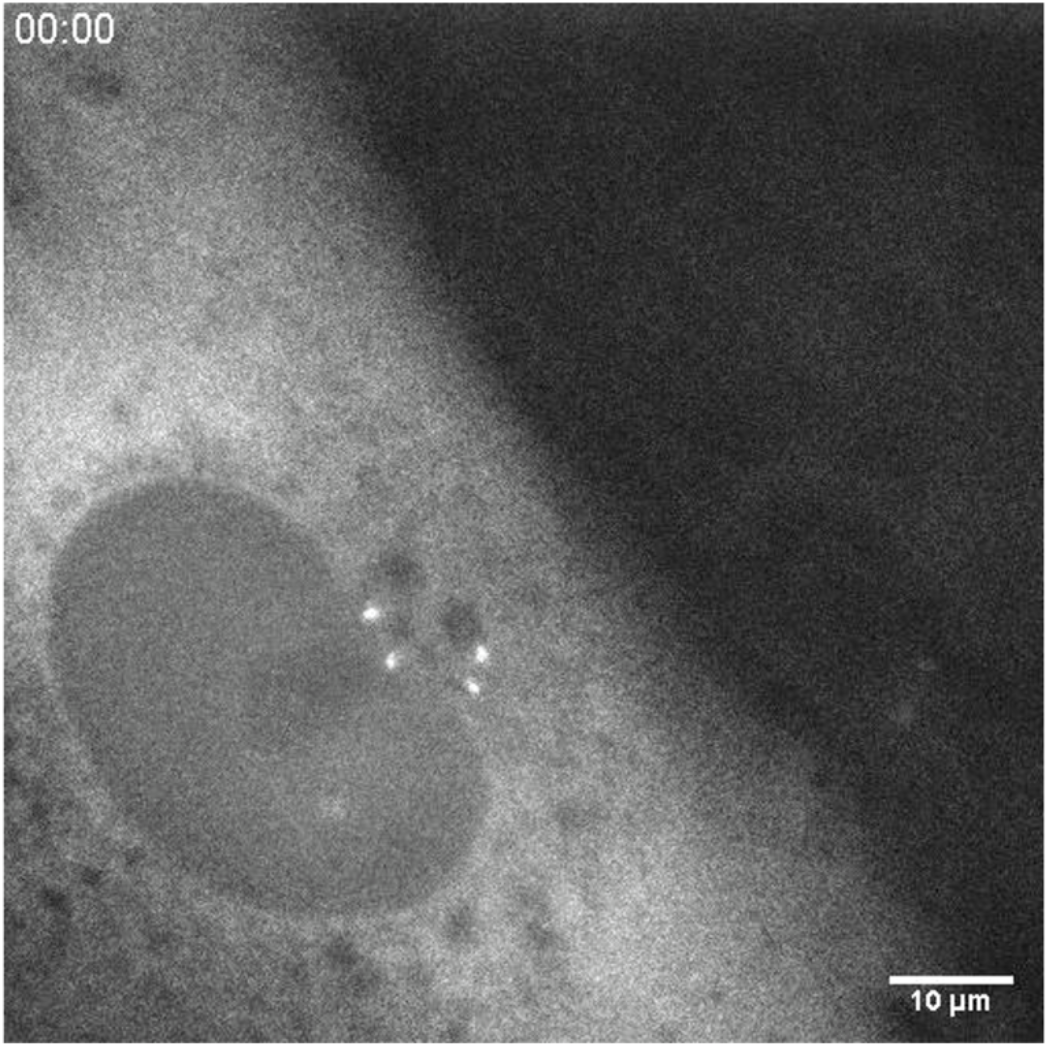
Partial supernumerary centriole disengagement in a differentiating TGC Time lapse movie showing a cell with supernumerary centrioles undergoing partial separation with subsequent pairs of centrioles; arrows show moments of separation. 10 minutes per frame, scale bar is 10μm.

**Supplementary Video 4:**
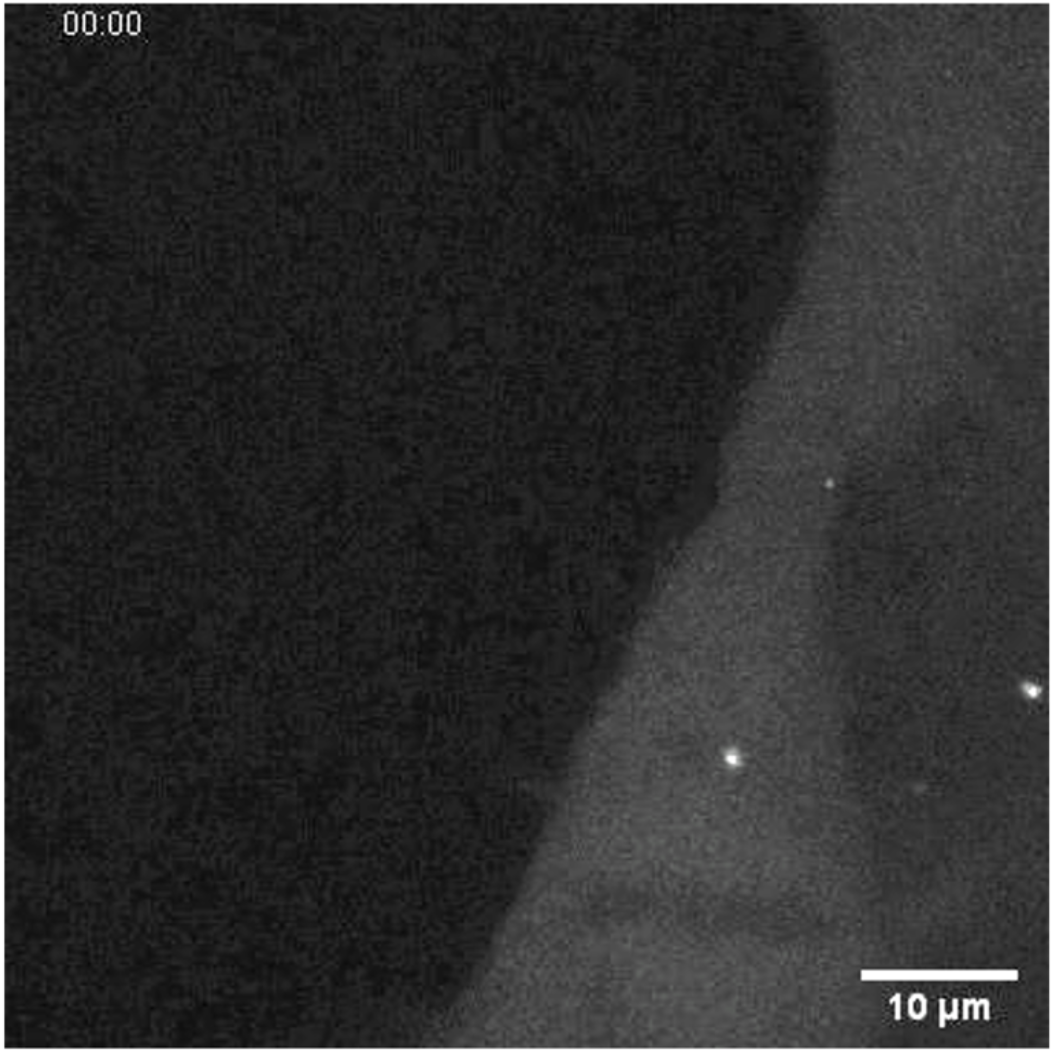
Post-mitotic centriole duplication and disengagement in a differentiating TGC Time lapse movie showing a cell that has just completed mitosis undergoing disengagement separation of mother and grandmother centrioles (first arrow) and their respective procentrioles (second and third arrows). 10 minutes per frame, scale bar is 10μm.

## Notes

### Competing Interest Statement

The authors have declared no competing interest.

